# Recent invasion of P transposable element into *Drosophila yakuba*

**DOI:** 10.1101/453829

**Authors:** Antonio Serrato-Capuchina, Stephania Zhang, Wendy Martin, David Peede, Eric Earley, Daniel R. Matute

**Affiliations:** Biology Department, University of North Carolina, Chapel Hill, N.C. USA

**Author notes:** Correspondence: Biology Department, University of North Carolina, Chapel Hill, North Carolina 250 Bell Tower Drive, Genome Sciences Building Chapel Hill, NC 27510, USA.

**Keywords:** P-element, hybrid dysgenesis, *Drosophila yakuba*, transposable elements

## Abstract

Transposable elements (TEs) are self-replicating genetic units that are common across prokaryotes and eukaryotes. They have been implicated in the origin of new molecular functions and in some cases, new phenotypes. Yet, the processes that lead to their evolution and how they enter the genome of their hosts remain largely underexplored. The P-element is one of the most well-known TEs in Eukaryotes, due to its rapid expansion in *Drosophila melanogaster* in the 1960s and its faster invasion of *D. simulans*, despite its fitness consequences in both species. Here, we describe a recent invasion of P-elements into *Drosophila yakuba*. Overall, PEs were found in *D. yakuba* with no PEs detected across its sister species, *D. teissieri* and *D. santomea*. These findings are surprising due the lack of a genetic bridge between *D. yakuba* and other *Drosophila* that harbor PEs, implicating a horizontal gene transfer mechanism similar to the one that gave rise to the invasion of PEs in *D. melanogaster* and *D. simulans*. We also report that the presence of these PEs causes a mild hybrid dysgenesis phenomenon; namely they cause a reduction in female reproductive potential (lower number of ovaries and ovarioles), but only at 29°C and not at 23°C. Given the ability of PEs to cross species boundaries and the fact that both *D. santomea* and *D. teissieri* have the ability to produce fertile progeny with *D. yakuba*, the *yakuba* species complex provides an opportunity to study PE spread through vertical transmission.

**ARTICLE SUMMARY:** P-elements (PEs) are transposons found in Neotropical *Drosophila* species. PEs have previously invaded two African *Drosophila* species where they rapidly increased in population frequency and fixed. We found that PEs invaded the genome of *D. yakuba*, an African species. In just 8 years, the frequency of the PEs increased from 0% to 18% but then decreased to 2%. This turnover shows that PE invasions can be transient. We found no evidence of full PEs *in D. yakuba*’ sister species, *D. santomea* and *D. teissieri*. PEs in this species complex can reveal the interplay between transposable elements and hybridization in nature.

## INTRODUCTION

Transposable elements (TEs) are autonomous genetic units that are able to propagate throughout the genome of a host (McClintock 1950, 1953). TEs are widespread across a vast range of organisms (reviewed in (Chuong *et al*. 2016)). In some cases, TE also prove to be a rapid method of genetic innovation (Werren 2011; Warren *et al*. 2015). TEs are often associated with the origin of new genetic and phenotypic diversity. In vertebrates, TEs have been shown to contribute to the evolution of gene circuits, leading to new lineage-specific gene regulation and functions. In the case of primates, for example, TEs can serve as a source of new variants in regulatory sequences (Trizzino *et al*. 2017). In angiosperms a significant portion of adaptive novelty is thought to be due to the activity of TEs (active TE-Thrust), resulting in gene duplications, novel expression patterns, and in some cases, gene disruptions (Debolt 2010; Ågren and Wright 2015). Other cases have shown the role of TEs in the origin of new phenotypes (Ding *et al*. 2016) and they have been commonly associated in the genetic basis of interspecific differences (Warren *et al*. 2015). TEs have often been associated with genome expansions across multiple taxa, potentially playing a role in the evolution of genome size (Vicient and Casacuberta 2017). In some of the most spectacular cases of genome expansions, fungal genomes are composed of up to 20% TEs, fish genomes are 55% TE, and in maize’ genome is up to 62% TEs (Sanmiguel and Bennetzen 1998; Daboussi and Capy 2003; Chalopin *et al*. 2015; Bilinski *et al*. 2018).

In the case of the *Drosophila*, TEs make up 20% of the genome (Eggleston *et al*. 1988) but this percentage varies depending on the population and species (Vieira *et al*. 1999). One of the best studied cases of the phenotypic effects of TEs in animals and across Eukaryotic systems — is P-elements (PEs) in *Drosophila*. PEs have rapidly spread worldwide throughout populations of the genetic model system, *D. melanogaster* (Engels and Preston 1980). Despite the self-replicating nature of TEs, this spread is puzzling, due to the negative phenotypic effects they cause. PEs in *D. melanogaster* lead to F1 sterility when the germ line of the female does not carry the molecular machinery regulating the expansion of PEs (Michalak 2009; Tasnim and Kelleher 2017). When a female who lacks PEs mates with a PE male carrier, the resulting F1s—both females and males — are sterile and show elevated rates of chromosomal breakage and increased mutation rates, a suite of traits referred to collectively as hybrid dysgenesis (HD) (Kidwell *et al*. 1977). Conversely, if a female with PEs mates with a PE male carrier, the F1s are fertile. In this case, the effects of PEs are silenced through a maternally inherited and germline-specific subclass of small non-coding RNAs, piRNAs (PIWI-interacting RNAs). This RNA facilitated silencing mechanism is not specific to PEs and has been shown to underlie repression throughout multiple classes of TEs and seems to play a role in repression across a variety of plant and mammalian hybrids (Michalak 2009). In *Drosophila*, F1 sterility due to the action of PEs is a simple and elegant model of how relatively simple genomic changes (i.e., the invasion of a TE) can induce reproductive isolation between genotypes rapidly and potentially lead to speciation (Serrato-Capuchina and Matute 2018).

PEs are thought to have originated in the *Drosophila willistoni* species group and, through a horizontal transfer event mediated by an unknown vector, invaded *D. melanogaster* (Kidwell 1992). In the case of *D. melanogaster*, despite the negative consequences associated with them, PEs managed to spread through vertical transmission throughout populations on every continent within 34 years (Kidwell 1983). PEs are also present in *D. simulans* but not in *D. sechellia*, two of the species that form the *simulans* clade, a sister group to *D. melanogaster*. PEs spread into *D. simulans’* entire range within 15 years (Kofler *et al*. 2015). Since the hybrid progeny between *D. melanogaster* and *D. simulans* are sterile (Sturtevant 1920; Ranz *et al*.2004), the invasion of PEs into *D. simulans* must have different origins outside of vertical transmission. As a result, a natural question is whether the genomes of related species have also been invaded by PEs and whether there are any conserved patterns in their transmission and phenotypic effects (Watson and Demuth 2013; Marco *et al*. 2018; Serrato-Capuchina and Matute 2018).

In order to understand how TEs regularly increase in frequency across a vast array of genomes it is crucial to study their spread, and potential reproductive tradeoffs, early in genomic invasions. For example, understanding the rate at which PEs increase in frequency through *Drosophila* species could inform how the HD phenomena arises and how PEs came to be so prevalent *D. melanogaster* and *D. simulans* populations. An ideal scenario to understand how PE increase in frequency is to obtain longitudinal samples and uncover an invasion in its early stages.

The *yakuba* species complex is an ideal model to study early TE expansion in natural populations. The complex consists of three species –*D. yakuba, D. santomea,* and *D. teissieri*– which exhibit diverse life history traits and wide geographic range. *Drosophila yakuba* is a widespread commensal species with similar life history traits to *D. melanogaster* and *D. simulans*. *Drosophila teissieri* is also widespread (although in fragmented populations, (Cobb *et al*. 2000) but is thought to be specialized to *Parinari* fruits (Lachaise *et al*. 1988; Comeault *et al*. 2017). The third species, *Drosophila santomea*, is endemic to the island of São Tomé where it is mainly found in undisturbed high montane forest (Lachaise *et al*. 2000; Matute 2010). Notably, the *yakuba* species complex has the only two known stable hybrid zones in the *melanogaster* species group. These zones exist between *D. yakuba* and *D. santomea* (Llopart *et al*. 2009; Turissini and Matute 2017), and between *D. yakuba* and *D. teissieri* (Cooper *et al*. 2017b).

We studied whether any of the species from the *yakuba* species complex harbored PEs and explored the phenotypic effects of recent PE invasions into an unexposed species. We performed a longitudinal study across 5 locations and 5 different time points in an attempt to pinpoint the early phenotypic effects of a PE invasion into novel species. We surveyed the *D. yakuba* clade across five collection years (ranging from 2003 to 2018) to explore whether the PE has spread into any of the three species. We found that *D. yakuba* harbors PEs, and that their frequency increased from 0% to 18% but then decreased to 2%. We did not find evidence of complete PEs in *D. yakuba’* sister species, *D. santomea* and *D. teissieri,* despite ecological overlap in geographic range. Additionally, we explored whether the hybrid dysgenic phenotype is expressed within *D. yakuba*, the species that recently acquired the PE. We tested for hybrid dysgenesis at two temperatures, 23° and 29°C, previously shown to be associated with HD in both *D. melanogaster* and *D. simulans* (Schaefer *et al*. 1979; Hill *et al*. 2016a). Our results show that PEs in *D. yakuba* can indeed cause HD but only at higher temperatures, which is consistent with the most deleterious effects of PEs in other *Drosophila* species. Notably, and unlike *D. melanogaster* and *D. simulans,* we found no HD at 23°C. Since no species that hybridizes with *D. yakuba* (and produce fertile progeny) carries functional PEs, the introduction of PEs is puzzling and suggestive of a horizontal gene transfer, mirroring invasions in other species of the *melanogaster* species subgroup. Our *D. yakuba* longitudinal sampling also revealed a sudden increase followed by a drastic drop in frequency of PEs which might shed light on the precise selective pressures that lead to rapid increases of this autonomous elements. Finally, our study presents the opportunity to study the dynamics of PE transmission across a hybridizing species complex, despite recently collected populations of *D. teissieri* and *D. santomea* not appearing to contain PEs.

## METHODS

### P-element detection from genome sequences

We used previously published genomes (Turissini and Matute 2017; Turissini *et al*. 2018) to assess whether there were PEs in the genomes of eight species of the *melanogaster* species group. Multiple methods have been proposed to quantify the allele frequency of known TE insertions in particular locations of the genome (reviewed in Serrato-Capuchina & Matute, 2018). However, our goal was to determine whether species from the *yakuba* clade harbor PEs anywhere in the genome. Therefore, we mapped raw reads from five *Drosophila* species to the *D. melanogaster* PE sequence (http://flybase.org/reports/FBte0000037.html) using bwa (Supplementary information) and calculated the number of mapped reads per million (rpm) (Kofler *et al*. 2018). We followed this procedure for the three species in the *yakuba* species group (*D. santomea*: *N*=17; *D. teissieri*: *N*=13; *D. yakuba*: *N=*109), as well as *D. simulans* (*N*=72), *D. sechellia* (N=XX), *D. mauritiana* (N=XX) and *D. melanogaster* (*N*=104). We also include reads from seven individuals of *D. orena* originated from the same isofemale line. The FASTQ file accession numbers of the sequences are listed in Table S1. Lines were considered to be candidates in harboring PEs if any read mapped to the PE sequence. For each species, we also calculated their mean log(rpm+1) and assessed whether that mean differed from zero (One Sample t-test, library stats, function ‘t.*test*’). We adjusted the critical P-values for significance to 0.01 to account for multiple comparisons (5 comparisons). To ensure comparable mapping results, we used Single End sequences only. For those lines that have been sequenced with Paired End reads, we used only one of the ends, chosen randomly.

### P-element detection at the population level

#### Fly lines

Our genomic survey revealed that species from the *D. yakuba* are likely to harbor PEs (See Results). Thus, we explored which individuals from a large collection of natural isolates from the three species in the group that harbored PEs. We collected females and males in the field from the three species across their geographical range (Table S2). We surveyed a total of 531 *D. yakuba*, 27 *D. teissieri*, and 336 *D. santomea* individuals. Populations were collected across 5 locations (Bioko, São Tomé, Príncipe, Cameroon, and Kenya; Figure S1) throughout their African range in 2003, 2009, 2013, 2015 and 2018. Collection details for each individual are listed in Table S2. 95% confidence intervals for point estimates of the proportion of individuals with PEs were calculated using the conjugate beta prior on the distribution of successes (library binom, function ‘*binom.cloglog*’ (Sundar Dorai-Raj and Sundar Dorai-Raj 2006)).

#### PCR

Using independent *yakuba* clade isofemale lines, we measured the frequency of PEs at different years in the three species of the *yakuba* species complex. To this end, we assessed whether individuals from these lines had any of the four exons that constitute a full PE. Since PEs require all four exons to be functional, our goal was to type all the individuals for each exon individually using PCR. We extracted genomic DNA from one female of each isoline (or an individual in ethanol) following the 96-well Puregene extraction kit protocol. To individually amplify each of the 4 exons that make up the full PE, we used primers described in (Hill, Schlötterer, & Betancourt, 2016). We did all PCRs using NEB reagents in a 10ul reaction (1ul 10x buffer, 1ul 10mM MgCl^2^, 0.5 ul 10mM dNTPs, 0.3 ul 10mM F+R primers, 1ul DNA, 0.05 Taq Polymerase, 5.85 ul H_2_0) with a thermocycling cycle of 92**°** denaturing, 59**°** annealing, 72**°** extension for 35 cycles in an Applied Biosystems 2720 Thermal Cycler. To score presence/absence of each exon, we ran 5ul of the PCR product in a 2% (APExBIO) agarose gel for 60 minutes at 120 volts and visualized the results using ethidium bromide staining. Sanger sequencing (Eurofins) was used to verify for PE presence in isolines that amplified for each primer to ensure the presence of the full continuous element.

#### Phylogenetic analysis

We aligned the sequence of each of the four exons of the PE found in *D. yakuba* with the sequence of their counterparts in *D. melanogaster*, *D. simulans*, *D. willistoni*, and *D. prosaltans* (accession numbers in Table S3) using MUSCLE (Edgar 2004). Exon sequences were limited to 500bp length primer amplifications (Kofler *et al*. 2015). Unrooted maximum likelihood trees were generated using RAxML with the transition/transversion ratio, proportion of invariant sites, and tree topology set to estimates. We calculated support for each branch by bootstrapping the tree 1,000 times. Trees were visualized with FigTree (Chevenet *et al*. 2006).

#### Hybrid dysgenesis

In crosses where PEs cause HD, F1 females and males from crosses between a PE^−^ females and a PE^+^ male (PE^−^/PE^+^) show stark gonadal defects. Females show atrophied ovaries and males show small testis (Raff *et al*. 1990). We explored whether *D. yakuba* F1s showed gonadal defects associated with HD. We scored four possible phenotypes *i*) presence of atrophied/rudimentary ovaries, *ii*) reduced number of ovarioles per ovary, *iii*) early onset of female reproductive senescence, and *iv*) reduced male fertility. We scored the four possible F1 genotypes following the steps as described below.

#### Crosses

We collected *D. yakuba* virgin flies from PE^+^ and PE^−^ isolines within 8 hours of eclosion and housed them in sex-specific vials. All flies were aged 4 to 9 days to minimize age effects. Crosses were performed by housing 5 individuals of each sex in a single vial. We made reciprocal F1s by crossing PE lines to non-PE carrying lines and produced the 4 types of possible progeny: PE^−^/PE^+^, PE^−^/PE^−^, PE^+^/PE^−^, PE^−^/PE^−^. To minimize the effect of different isofemale lines, we used a random number generator to determine the isolines that were mated. In total, we used 5 isolines of PE containing São Tomé *D. yakuba* and 5 isolines of non-PE containing *D. yakuba*. We left the vials undisturbed at their test temperatures until removing the adults after 5 days, adding a 0.05% propionic acid solution and a pupation substrate to the food (Kimwipe, Kimberly Clark). F1s were collected as virgins daily and separated by sex. All crosses were performed both at 23°C and at 29°C, to measure the effect of temperature in the magnitude of HD.

#### Gonad number—Counts

First, we scored whether F1 had 0,1 or 2 developed gonads, with healthy females and males having 2 ovaries and 2 testes respectively, at both 23° and 29°C. After 4 to 9 days, flies were anesthetized with CO_2_ and their gonads removed with metallic forceps (Wong and Schedl 2006). Gonads from each individual were subsequently fixed on a precleaned glass slide with chilled *Drosophila* Ringer’s solution (Cold Spring Harbor Protocols). We counted the number of non-atrophied gonads for each individual. Ovaries were considered atrophied if they had no ovarioles. Testes were considered atrophied if they had less than half the length of wild-type testes, however, all males contained wild-type testes. In the case of females, we also counted the number of ovarioles (see below) in each mature ovary using a Leica, S6E stereoscopic microscope. We scored 273 females at 23°C and 186 females at 29°C. Table 1 shows the number of females dissected for each genotype. For ovariole counts, we only scored flies for which the dissection contained both left and right gonads.

**TABLE 1.**
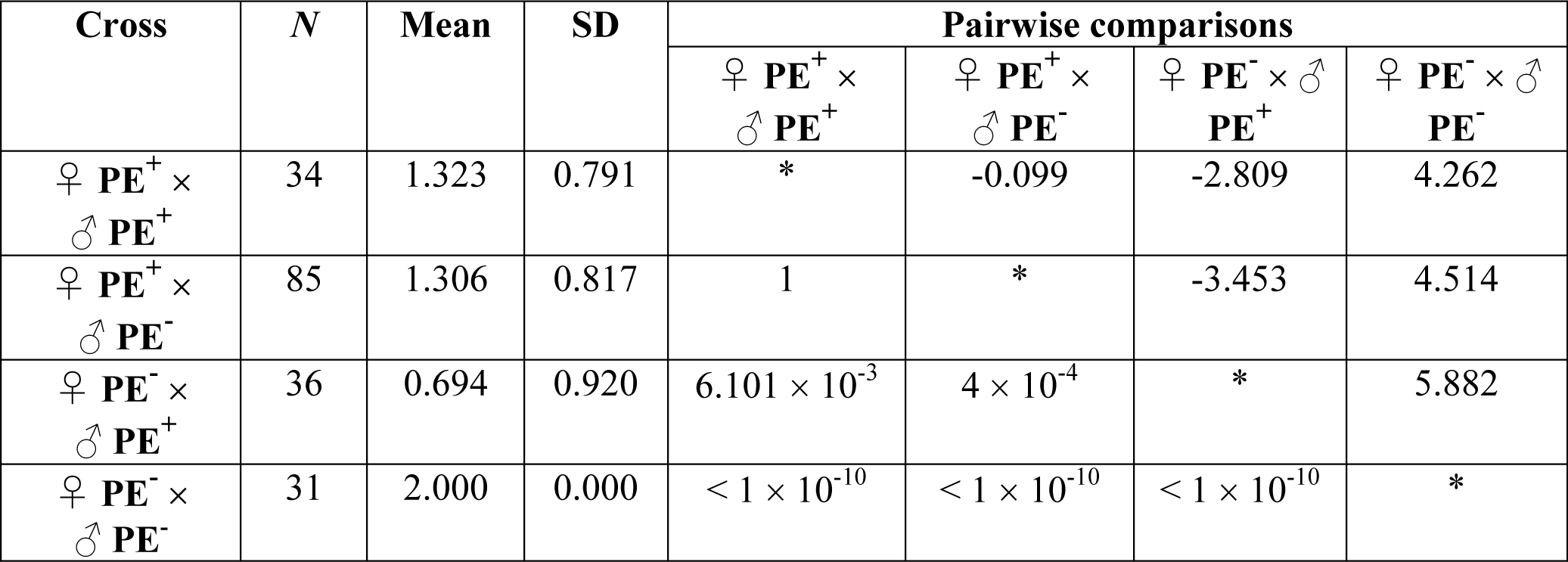
The presence of PEs affects the number of ovaries in F1 *D. yakuba* female (from intraspecific matings) at 29°C. *N* is the number of dissected females that produced the means (percentage of females mated) and standard deviations (SD). The last four columns show pairwise comparisons as 4 × 4 matrices for each cross. The upper triangular matrix shows the Z value from an approximate Two-Sample Fisher-Pitman Permutation Test (9,999 permutations). The lower triangular matrix shows the P-value associated to the comparison. Only pairwise comparisons with P < 0.008 were considered significant.

#### Ovary number—Statistical analyses

We scored whether each F1 female had 0, 1, or 2 ovaries as described above. To quantify the magnitude of heterogeneity among F1 genotypes, we fitted a multinomial regression using the function *multinom* in the library *nnet* (Venables *et al*. 2003) where the number of ovaries was the response of the multinomial assay and the mother and father genotypes were the fixed effects. We also included the interaction between these two effects to account for the interplay between the genome of the two parents. The significance of the effects was inferred using the function *set_sum_contrasts* (library *car* (Fox and Sanford 2011)), and a type III ANOVA (library *stats* (R-Core-Team 2013)) in R. Since we did experiments at two different temperatures (23°C and 29°C), we fitted two multinomial regressions. To do post-hoc comparisons between crosses, we used a Two-Sample Fisher-Pitman Permutation Test (library *coin,* function ‘*oneway_test’*; (Hothorn *et al*. 2006)) and adjusted the critical P-values for significance to 0.008 to account for multiple comparisons (6 comparisons).

#### Ovariole number—statistical analyses

A second potential phenotype of HD is the reduction in the number of ovarioles per ovary in female F1s, in females that did not show atrophied ovaries (Khurana *et al*. 2011). In these females, even with 2 ovaries, their reproductive potential can be limited through a lack of ovarioles (Lobell *et al*. 2017). We quantified whether the genotype of the mother, of the father, or the interaction between these two terms affected the number of ovarioles. We analyzed the mean number of ovarioles per ovary (i.e., females with two ovaries will have more total ovarioles than females with one ovary) to account for difference in the number of ovaries. We excluded those females that showed completely atrophied ovarioles from this analysis. We used a Poisson-distributed linear model (library *stats,* function ‘*glm’* (R-Core-Team 2013)). To assess the significance of interactions, we followed a maximum-likelihood model simplification approach (Crawley 1993); we first fitted a fully factorial model containing all factors and interactions and then simplified it by a series of stepwise comparisons, starting with the highest-order interaction and progressing to lower-order interaction terms and then to main effects.

#### Female reproductive senescence—counts

We explored whether the age of the female had an effect on the number of ovarioles in PE^+^ and PE^−^ females. Specifically, we explored whether HD manifested itself as a shorter reproductive period in females that carried PEs (Lobell *et al*. 2017). In this scenario PE*^+^* will show a sharper decline in their ovariole number compared to their PE^−^ females. To score females of different age, we cleared bottles and collected newly eclosed virgins within 8 hours of clearing as described above (Section ‘*Crosses’*). To account for heterogeneity across lines, we studied 5 different isolines per population type: 5 PE^+^ isofemale lines from São Tomé, 5 PE^−^ isofemale lines from São Tomé, and 5 PE^−^ isofemale lines from the African continent, for a total of 15 isofemale lines per time point. Female virgins were then dissected every 5 days for 25 days to count the ovariole count as they aged. In total there were 1,125 observations: 5 time points × 5 isolines × 15 individuals per line × 3 distinct population types.

#### Female reproductive senescence—Statistical analyses

We used an Analysis of covariance (ANCOVA) to assess whether the presence of PEs affected the reproductive capacity of a female at different ages. We used the function *lm* in the R library *stats* (R-Core-Team 2013). First, we used the regression coefficients from the ANCOVA to compare the intercept of the linear regressions of females with and without PEs. This test assessed whether genotypes had inherent differences in the number of ovarioles (i.e., whether the effect of genotype—if a female is PE^+^ or PE^−^—was significant). Second, we compared the rate of decline of fertility among genotypes. To this end, we quantified differences in the slope of the regressions of number of ovarioles as age progressed (i.e., the interaction between female age and her genotype). To evaluate the significance of the interaction, we used information obtained with the function *lm* as described immediately above and also performed a likelihood ratio test (LRT; function *lrtest*, R library *lmtest* (Kuznetsova *et al*. 2015)).

#### Male fertility—sperm motility

We scored whether F1 male progeny produced motile sperm. We dissected the testes of each individual with metallic forceps (Miltex Catalogue number: 17-301, McKesson, Richmond, VA) and mounted them on chilled Ringer’s solution. We mounted up to five males per slide and scored whether they had motile sperm within 5 minutes of starting the first dissection. We scored 843 F1 males at 23°C and 542 F1 males at 29°C. To quantify the effect of the genotype on sperm motility among F1 genotypes, we fitted a binomial regression (library stats, function ‘*glm’*). Whether a male had fertile sperm or not was the response of the binomial model, while the mother and father genotypes were the fixed effects. We also included the interaction between these two effects to account for the interplay between the genome of the two parents. We used LRTs (described above) to test whether to retain the interaction and the fixed effects. We found no sterile males at 23°C, so we only fitted a single linear model at 29°C. To do posthoc tests, we used a Tukey Honest significance difference test (library multcomp, function ‘*glht’*).

#### Male fertility— progeny count

Finally, we scored whether F1 male progeny showed reduced fertility despite showing normal size testes (see above). We collected F1 males from the four F1 genotypes raised at the two studied temperatures (23°C and 29°C) and mated them to virgin PE-females. We watched the matings to ensure they were not abnormally short (less than 10 minutes (Matute and Coyne 2010)); as soon as the mating was over, we removed the male from the vial. We let the female lay eggs for 10 days. After this period, we removed the females and let the progeny develop at 23°C. Every two days, we counted the progeny produced by each female until no more flies emerged. We quantified the heterogeneity of the amount of progeny using a generalized linear model similar to the one described above (section ‘Ovariole number—statistical analyses’) where the number of progeny produced by each individual female was the response, the genotype of the cross and temperature at which the cross was performed were the fixed effects.

## Data availability

Supplemental Material, File S1 contains supplementary figures, supplementary tables and supplementary legends. Figures S1 and S2 are supplementary figures. Table S1 lists all the short read accessions used in this manuscript. Table S2 lists the flies screened with PCR. Isofemale lines are available upon request. Table S3 lists this PE sequence accession numbers. Tables S4-S6 report supplementary results.

The code used for all analyses reported here is available on Dryad (Accession number: TBD). All counts, raw pictures, and datasets are also deposited in Dryad.

## RESULTS

### Genome wide detection of P-elements in *D. yakuba*, *D. santomea* and *D. teissieri*

We used previously published genome sequences for isofemale lines in the *D. melanogaster* species group to test for the presence of the PE sequence. The provenance of these genomes is geographically heterogeneous and includes lines from multiple locations (Table S1). Our analyses included eight of the nine species of the *melanogaster* species subgroup (*D. yakuba, D. teissieri, D. santomea, D. mauritiana, D. sechellia, D. orena, D. simulans* and *D. melanogaster*). The last two have previously been reported to harbor PEs. All tested species showed at least one read mapping to a portion of the PE sequence, of which the D. *melanogaster* and *D. simulans* genomes showed the highest signal for presence of PEs, followed by *D. yakuba* (Figure 1). (*D. melanogaster* is not shown as all 104 lines had reads mapping to the PE sequence).

**FIGURE 1.**
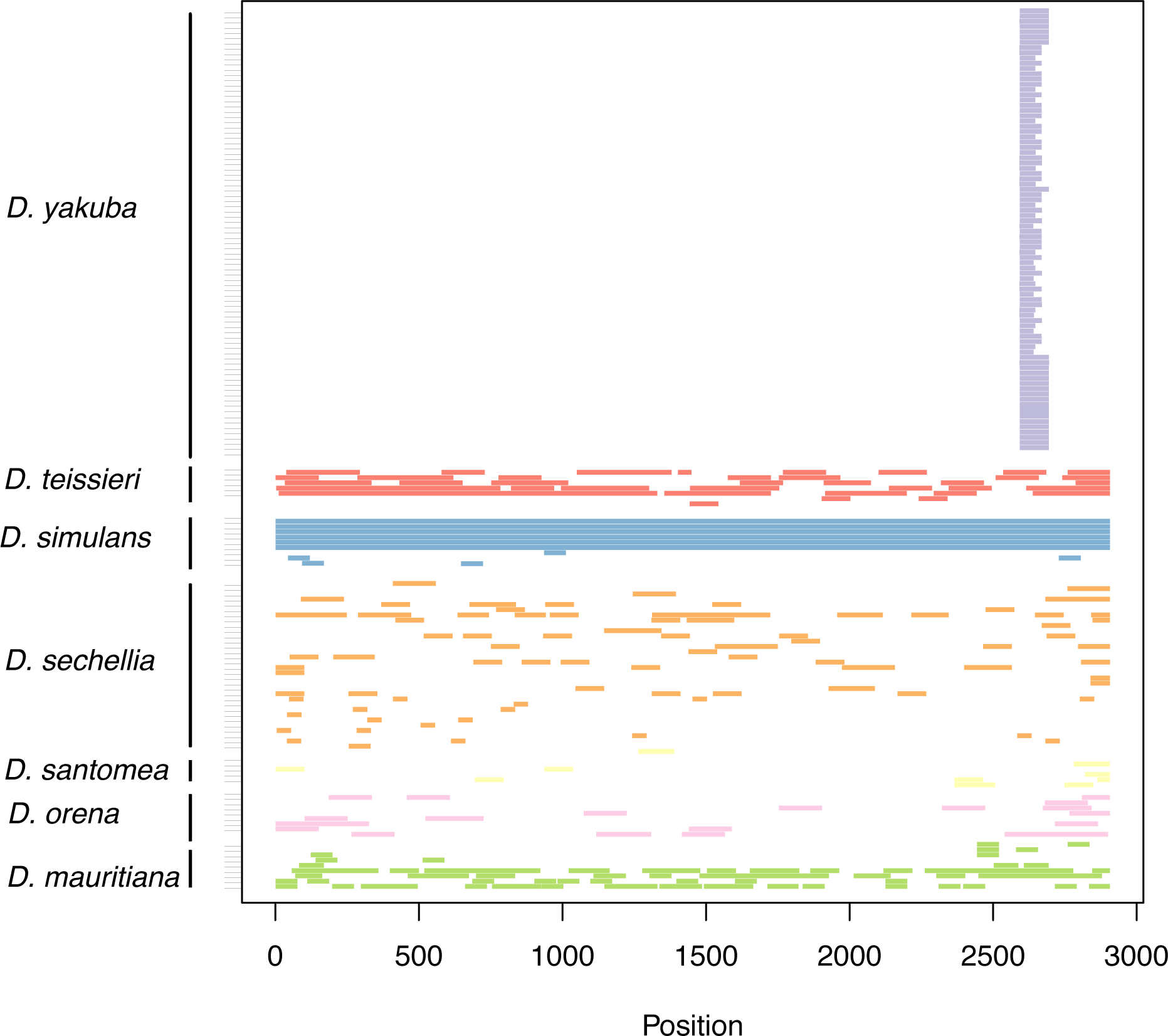
Genome sequences suggest P-elements might be present in the multiple species of the *melanogaster* species group. Each row represents the genome of an individual isofemale line or fly. Each colored block shows reads that mapped to the PE sequence of *D. melanogaster*.

Figure S2 show the log(rpm+1) for the *yakuba* species clade. There is extensive variation in the number of reads that map to the PE sequence across species (One-way ANOVA, F_4,310_ = 872.32, P < 1 × 10^−10^). This heterogeneity persists even after only the lines from the *yakuba* clade are included (F_2,136_=7.454, P= 8.474 × 10^−4^), mainly driven by the fact that all *D. santomea* lines showed low coverage for the PEs. These mapping results are consistent with previous reports of *D. melanogaster* and *D. simulans* lines that contain PEs (Kofler *et al*. 2015) and gave rise to the hypothesis that other species in the *melanogaster* species subgroup might harbor PEs. We tested this hypothesis for each of the species in the *yakuba* clade using PCR by amplifying each PE exon individually.

### P-elements have changed in frequency in *D. yakuba* from São Tomé

Next, we focused on species from the *D*. *yakuba* species group and scored the proportion of individuals that harbor a PE, across multiple years, throughout continental and island populations. This approach allowed us a temporal and longitudinal snapshot of PE spread within each species of the *yakuba* clade; *D. yakuba*, *D. teissieri,* and *D. santomea*. The results from the each of three species within the *D. yakuba* species complex are described as follows.

#### Drosophila yakuba

PCR amplification showed no evidence for PEs across continental lines (lines scored in Table S2). Similarly, before 2013, *D. yakuba* collections from São Tomé, showed no evidence of PEs. Additionally, we found no PEs in individuals collected in the island of Bioko, 460kms to the north of São Tomé. As of 2015, 18% of the *D. yakuba* individuals collected in São Tomé harbored PEs (19 out of 106 individuals, 95% confidence interval: [11.32%−25.76%]), with the frequency decreasing to 2% in 2018 (4 out of 200, 95% confidence interval: [0.08%−4.65%]; Figure 2). It is worth noting that 61 out of 66 lines whose genome was sequenced (all collected before 2010) showed at least one read that mapped to the PE sequence, albeit at low levels. Notably, all reads map to a single terminal region in Exon 3 (Figure 1) suggesting that none of these lines had a full PE.

**FIGURE 2.**
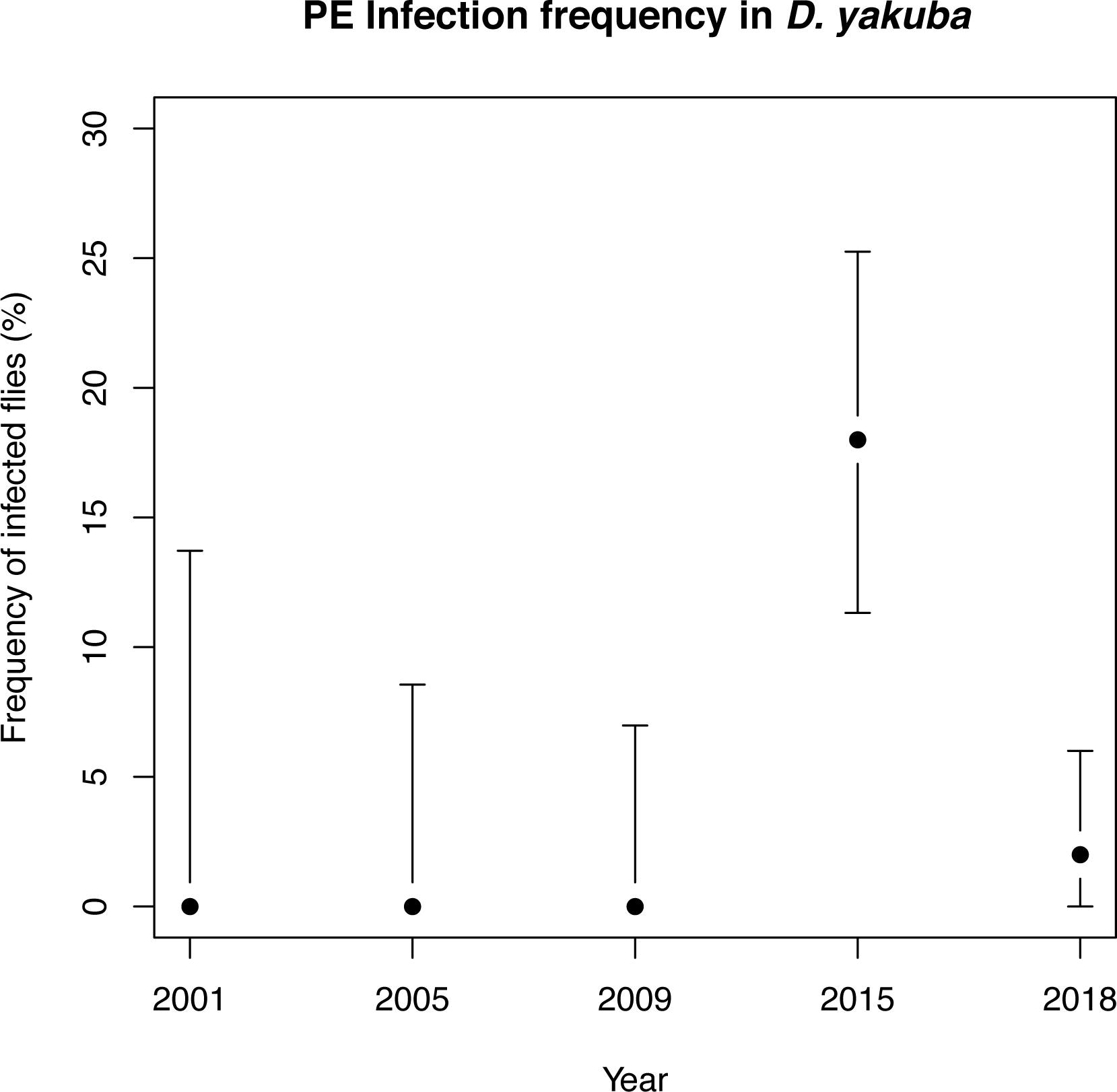
Frequency of the PE is *D. yakuba* in collections from five different years in the island of São Tomé. Frequency represents the proportion of individuals that show evidence for any of the four exons of the PE in a PCR test. The black dot represents the actual measured frequency and the bars show the 95% confidence intervals calculated as Bayesian binomial intervals. PE Infection frequency in *D. yakuba*

We assessed whether individuals from these populations had the full PE. In 2015, of the 19 individuals that were PE positive, only 6 of them contained the entire PE, with the remaining 13 lacking at least 1 of the 4 exons. In 2018, of the four individuals that were PE positive, two contained the full element and the other two contained partial elements. The detailed results of this screening are shown in Table S4. In the span of 2 years (2013 to 2015) the proportion of individual *D. yakuba* in São Tomé with PEs increased from 0% to 18% but by 2018 PEs were found in only 2% of lines (N=200) sampled on the same island.

#### Drosophila teissieri

*D. teissieri* is present across continental Africa and its neighboring island of Bioko. We examined 27 lines collected between 1970 and 2015. The majority of the lines were collected in the island of Bioko between 2009 and 2013. We found only one individual that contained any portion of the PE, a typed female from the isofemale TBRAZ28 (collected in Brazzaville, Republic of Congo), which contained exons 0 and 3 of the P-element. Since a functional PE requires all the four exons, we conclude that either no functional P-element is present in any *D. teissieri* line or that PEs are segregating at a population frequency lower than 1/27. This result differs from our genome detection approach but are not inconsistent. The lines Cascade 4.3, Cascade 4.1, and Cascade 2.4, all from the island of Bioko show no PCR amplicons for any of the PE exons but show reads that map to the PE sequence. The short read coverage does not suggest the presence of a continuous (and active) PE. Some of the missing sequence is precisely where the PCR primers anneal, thus explaining why the PCR scans did not detect them.

#### D. santomea

Lastly, we scored *D. santomea,* the sister species of *D. yakuba*. Since this species is endemic to the island of São Tomé, all the lines we studied were collected in this island (N=236 lines). We found no evidence for any of the four exons of the PE, strongly suggesting that either the PE is not present in this species or that it segregates at a population frequency lower than (1/336).

### P-element genealogy

We built a phylogeny using the sequence of the PEs found in the *melanogaster* species subgroup (*D. melanogaster*, *D. simulans* and *D. yakuba*). We found that the PE sequences are not partitioned by species (Figure 3). This result is consistent with the longitudinal sampling in these three species which suggests recent invasions of the PEs. The invasion seems to be recent enough that none of the PEs have accumulated any distinct differences.

**FIGURE 3.**
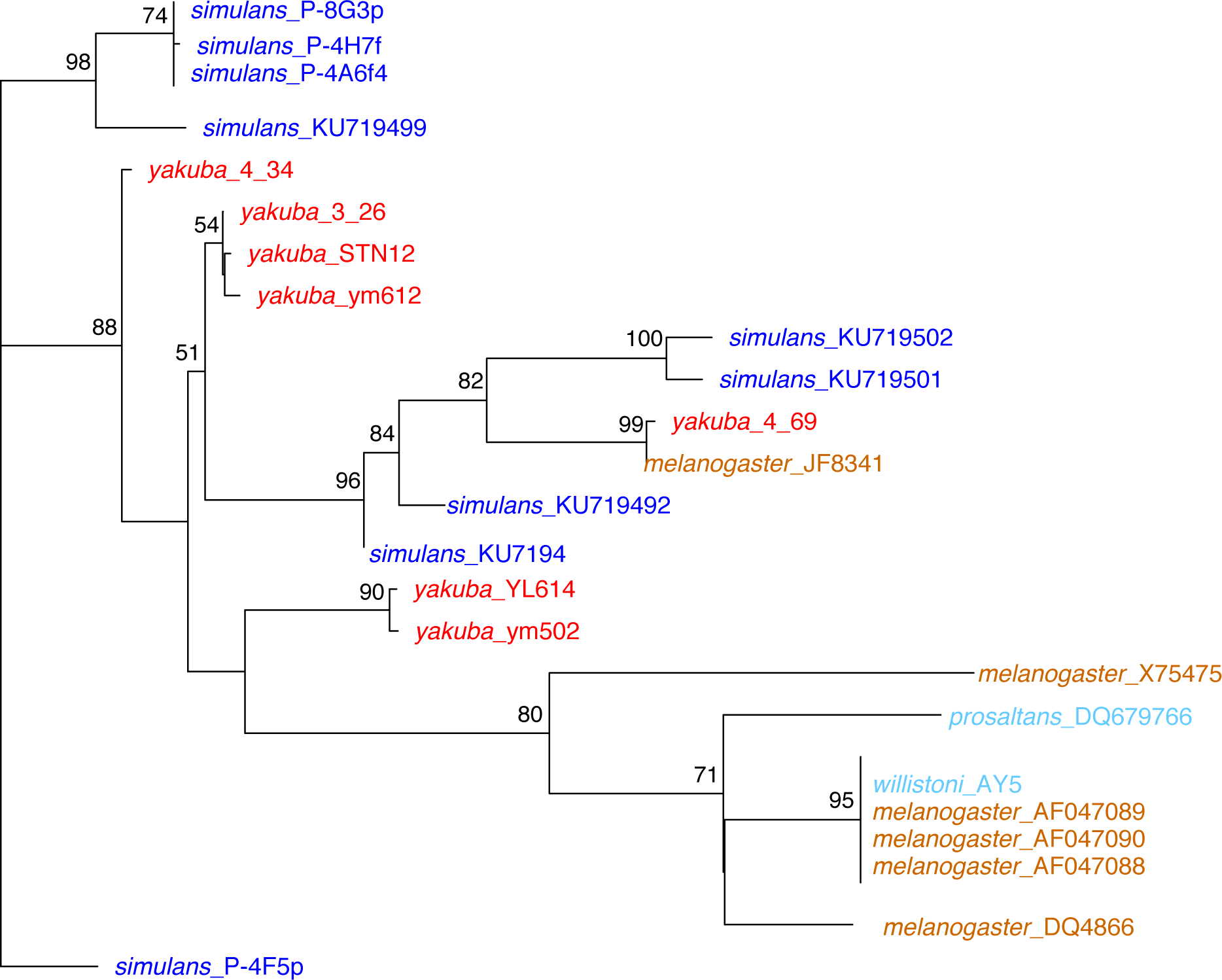
P-element sequences found in five species of *Drosophila* are not partitioned by species. A maximum likelihood unrooted tree indicates that the P-elements in different species of *Drosophila* have not accumulated species-specific mutations suggesting recent horizontal gene transfer. Number above nodes convey bootstrap support (1,000 replicates). Bootstrap values below 50% are not shown.

### P-elements cause mild hybrid dysgenesis in *D. yakuba*

Since the presence of PEs is polymorphic across *D. yakuba*, a natural hypothesis is that their presence might cause hybrid dysgenesis in PE^−^/PE^+^ individuals, mirroring effects described in both *D. melanogaster* and *D. simulans*. We assessed four possible outcomes of PEs in isolines and their resulting F1s consistent with the described effects of hybrid dysgenesis in other species: reduced number of ovaries, reduced number of ovarioles per ovary in females, early onset of reproductive senescence, and lack of sperm/reduced fertility in males. We describe each of these phenotypes as follows.

#### Ovary number

First, we scored whether females from the four possible genotypes (♀PE^+^/♂PE^+^, ♀PE^+^/♂PE^−^, ♀PE^−^/♂PE^+^, and ♀PE^−^/♂PE^−^) had 0, 1, or 2 ovaries. Wild type females usually have 2, while dysgenic flies show either 0 or 1 ovary (Engels and Preston 1980). We dissected females produced at two different temperatures, 23°C and 29°C. At 23°C, we found no heterogeneity in the number of ovaries among parental or F1 female genotypes as every single female (*N* > 21 per genotype) had 2 ovaries (as observed in Figure 4A).

**FIGURE 4.**
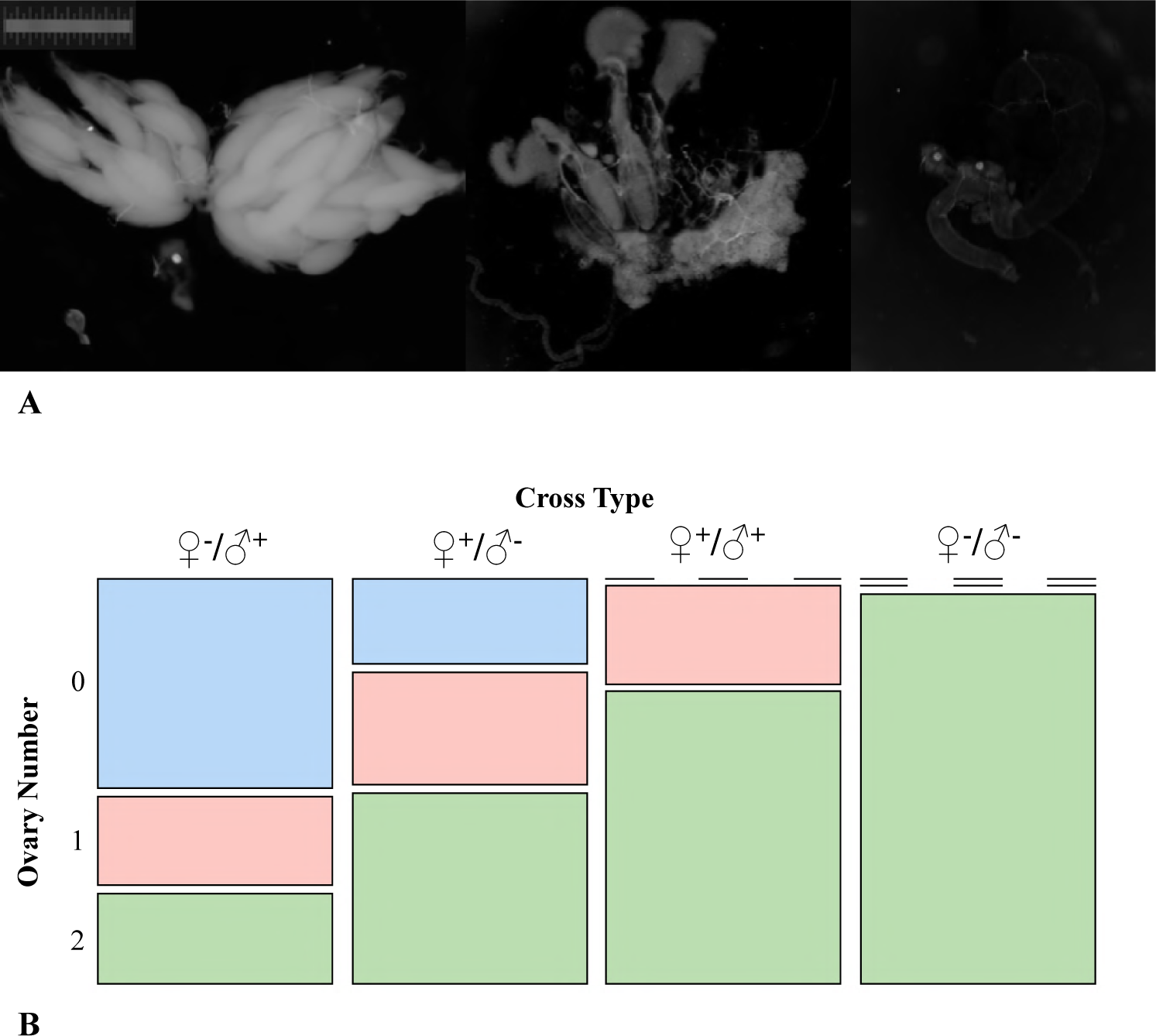
Hybrid dysgenesis in *D. yakuba* in the form of atrophied ovaries at 29°C. A: All females shown in this figure have genotype PE^+^/PE^+^ and were raised at 29°C. <u>Left:</u> Female with two ovaries. All F1s raised at 23°C, regardless of their genotype, also have this phenotype. <u>Middle</u>: Female with one functional (albeit reduced) and one completely atrophied ovary. <u>Right</u>: Female with two atrophied ovaries. Females in this latter category have no ovarioles. (Scale: Total length = 1mm, each division = 0.01mm). **B:** Mosaic plot of the proportional number of ovaries per cross type (designated female/male and PE status) at 29°C. F1 ovary number at 23°C is not shown as all the scored females, regardless of their genotype, have 2 ovaries.

However, the number of ovaries between genotypes differs at 29°C. Among F1 females, we found multiple individuals with no ovaries (Figure 4B) and significant heterogeneity in the number of ovaries by type of cross when reared at 29°C (Figure 4B). The mean, standard deviations, and post-hoc pairwise comparisons (permutation based) are shown in Table 1. ♀PE^−^/♂PE^−^ F1 females always have two ovaries (N=21). All the other three F1 genotypes showed fewer ovaries than the ♀PE^−^/♂PE^−^ cross (Table 1, Figure 4). F1s resulting from crosses in which only the male contained the full PE (♀PE^−^/♂PE^+^) resulted in the most severe fitness costs with 50% of F1s containing no ovaries (Table 1, Figure 4). In F1s resulting from crosses in which only the female contained the full PE (♀PE^+^/♂PE^−^), the number of F1s without ovaries was close to 15% (Figures 2 and 3A, Table 1). Finally, in cases where both parentals contained full PEs, no F1s had a complete loss of ovaries, with 15% of F1s containing 1 ovary and the remainder containing both ovaries (Figures 3 and 4A, Table 1). The heterogeneity in ovary number was due the interaction of the mother and the father genotype (LR=29.7097, df=6, P=4.463 ×10^-5^). These results are consistent with the hypothesis that PEs cause a HD syndrome where ♀PE^−^/♂PE^+^ F1s are the most affected, a phenomenon similar to the one observed in *D. simulans* and *D. melanogaster*.

#### Ovariole number

Hybrid dysgenesis can manifest itself not only as the absence of ovaries but also through the development of “rudimentary” ovaries, i.e. ovaries with fewer ovarioles. We counted the mean number of ovarioles in F1 females in the four types of F1 progeny. For these analyses, we excluded all individuals that had completely atrophied ovaries and consequently no ovarioles. At 23°C we found heterogeneity across genotypes and the source of that variation was the interaction between the female and the male genotype (Figure 5A, Table S5). Surprisingly, ♀PE^+^/♂PE^−^ F1 female progeny had more ovarioles than the mean from any other cross direction (an average of a 48% increase, Figure 5B). All other crosses produced an average of 25 total ovarioles and were no different from each other (Table 2). This result suggests, that at least in *D. yakuba*, females with PEs might show the conditional fitness advantage of increased fertility when mated to males that do not have PEs.

**FIGURE 5.**
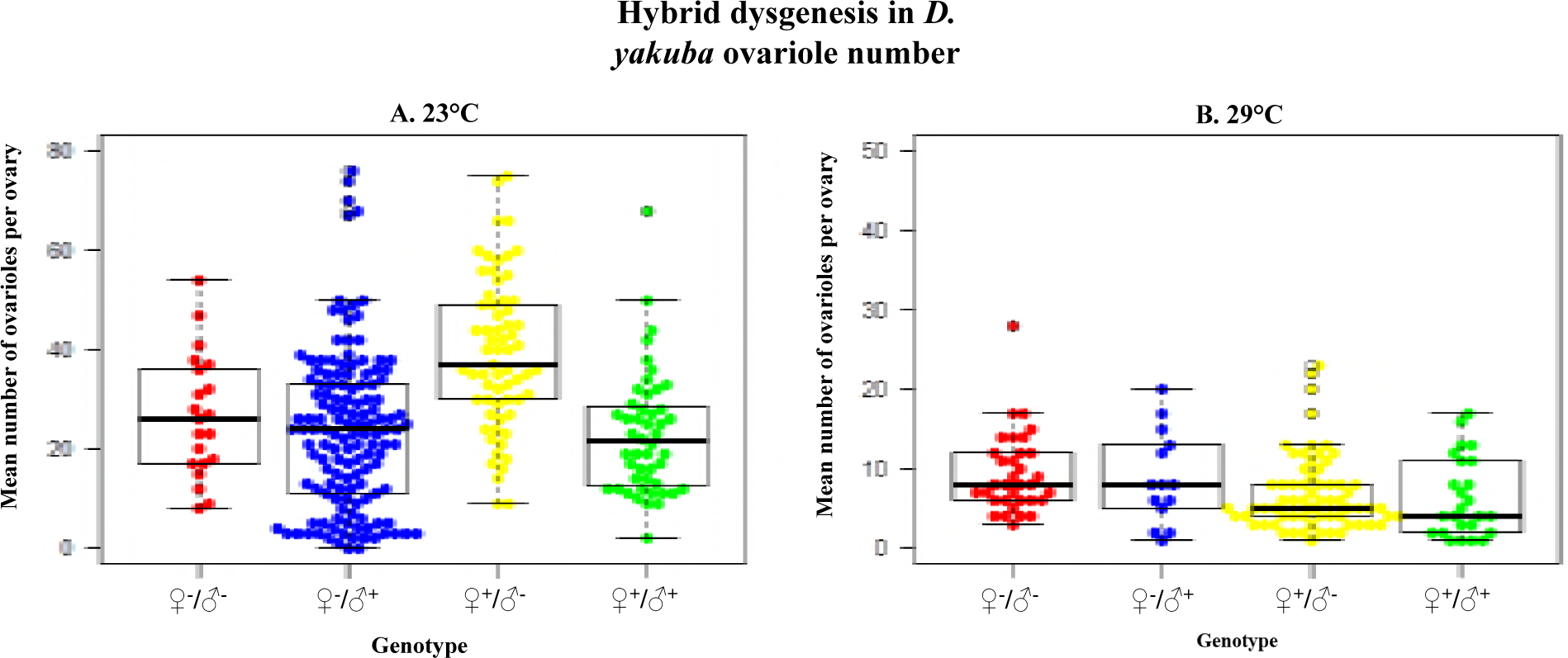
Hybrid dysgenesis in *D. yakuba* in the form of reduced number of ovarioles per ovary at 29°C. The boxplots show the ovariole number in the four possible F1 genotypes at two different temperatures: 23°C **(A)** and 29°C **(B).**

**TABLE 2.** The presence of PEs affects the number of ovarioles in F1 *D. yakuba* females (from intraspecific matings) at 23°C and at 29°C. *N* is the number of dissected females that produced the means (percentage of females mated) and standard deviations (SD). The last four columns show pairwise comparisons as 4 × 4 matrices for each cross. The upper triangular matrix shows the Z value from an approximate Two-Sample Fisher-Pitman Permutation Test (9,999 permutations). The lower triangular matrix shows the P-value associated to the comparison. Only pairwise comparisons with P < 0.008 were considered significant.

| 23°C |  |  |  |  |  |  |  |
| --- | --- | --- | --- | --- | --- | --- | --- |
| Cross | *N* | Mean | SD | Pairwise comparisons |  |  |  |
|  |  |  |  | ♀ PE<sup>+</sup> × ♂ PE<sup>+</sup> | ♀ PE<sup>+</sup> × ♂ PE<sup>-</sup> | ♀ PE<sup>-</sup> × ♂ PE<sup>+</sup> | ♀ PE<sup>-</sup> × ♂ PE<sup>-</sup> |
| ♀ PE<sup>+</sup> × ♂ PE<sup>+</sup> | 49 | 11.469 | 6.252 | * | 5.4109 | 0.492 | 1.286 |
| ♀ PE<sup>+</sup> × ♂ PE<sup>-</sup> | 65 | 19.800 | 7.571 | < 1 × 10<sup>-10</sup> | * | -5.947 | -3.218 |
| ♀ PE<sup>-</sup> × ♂ PE<sup>+</sup> | 138 | 12.089 | 7.978 | 0.629 | < 1 × 10<sup>-10</sup> | * | 0.816 |
| ♀ PE<sup>-</sup> × ♂ PE<sup>-</sup> | 21 | 13.571 | 6.201 | 0.206 | 7.001 × 10<sup>-4</sup> | 0.411 | * |
| 29°C |  |  |  |  |  |  |  |
| Cross | *N* | Mean | SD | Pairwise comparisons |  |  |  |
|  |  |  |  | ♀ PE<sup>+</sup> × ♂ PE<sup>+</sup> | ♀ PE<sup>+</sup> × ♂ PE<sup>-</sup> | ♀ PE<sup>-</sup> × ♂ PE<sup>+</sup> | ♀ PE<sup>-</sup> × ♂ PE<sup>-</sup> |
| ♀ PE<sup>+</sup> × ♂ PE<sup>+</sup> | 15 | 6.360 | 4.974 | * | -0.1676 | -1.433 | 2.814 |
| ♀ PE<sup>+</sup> × ♂ PE<sup>-</sup> | 66 | 6.909 | 4.748 | 0.874 | * | 3.724 | 3.724 |
| ♀ PE<sup>-</sup> × ♂ PE<sup>+</sup> | 14 | 8.786 | 5.847 | 0.158 | < 1 × 10<sup>-10</sup> | * | 4.024 |
| ♀ PE<sup>-</sup> × ♂ PE<sup>-</sup> | 34 | 9.382 | 5.069 | 0.005 | 1 × 10<sup>-4</sup> | < 1 × 10<sup>-10</sup> | * |

Generally, at 29°C, independent of the male used in cross, F1s produced from PE^−^females have more ovarioles than PE+ females. ♀PE^−^/♂PE^+^ females have fewer ovarioles than ♀PE^−^/♂PE^−^ females (Z-test on regression coefficients = 4.477, P= 7.58 ×10^-6^; Table S5). However, we found no difference in the number of ovarioles between ♀PE^−^/♂PE^+^ and ♀PE^+^/♂ PE^+^ females (Table 2). The pairwise comparisons (shown in Table 2) indicate that the only clear pattern is that ♀PE^−^/♂PE^−^ F1s have more ovarioles than females from any of the other three genotypes. It is worth noting that our power to detect differences at 29°C is lower as all the genotypes show lower fertility than at 23°C. Just as it occurs with ovary number, the PE-induced reduction in ovariole number is temperature dependent and only occurs at high temperature.

#### Reproductive senescence

A third potential phenotype in hybrid dysgenesis is that PE-carrying females show a rapid decrease in fertility as they age (Schnebel and Grossfield 1988). Specifically, we tested whether the presence of PEs was predictive of reproductive output throughout the lifespan of females. We tested this possibility by counting the number of ovarioles of females with and without PEs at five different ages for 25 days (Figure 6). As expected (Wayne *et al*. 2006), the number of ovarioles decreases as females age (Table 3). The intercept was similar for both types of females which indicates the initial reproductive potential is similar in females carrying and not carrying PEs (genotype effect: Table 3). Additionally, the rate of decrease (i.e., the slope of the linear regression) was not different for the two regressions either (genotype by age interaction: Table 3). These results indicate that, at least at 23°C, PEs in *D. yakuba* do not induce early reproductive senescence.

**FIGURE 6.**
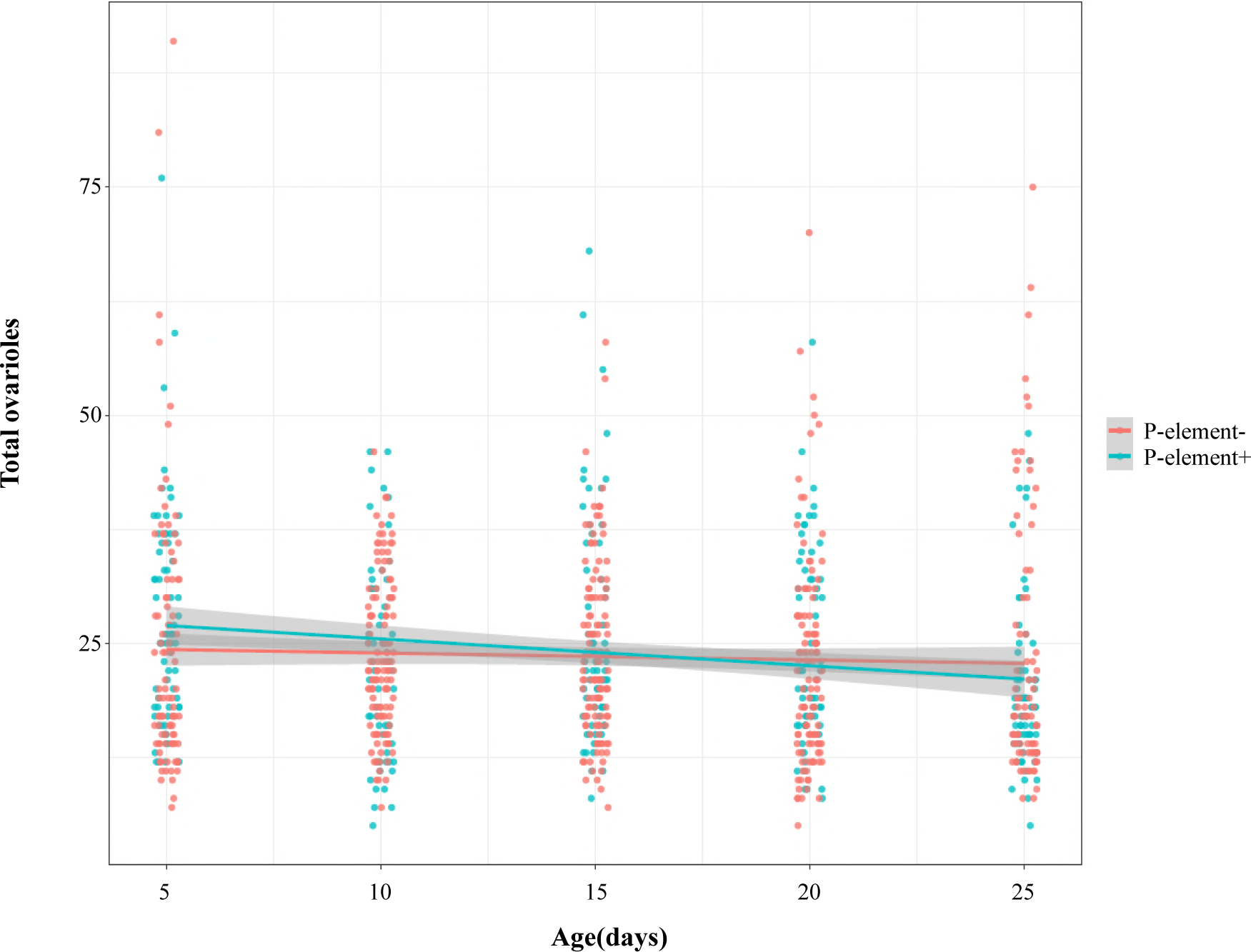
Number of ovarioles observed in PE+ and PE‐ females as they age. Red points show the observations for the ten PE^−^ lines. The red line shows the linear regression for these observations. Blue dots show the observations for the five PE^+^ lines. The blue line shows the linear regression for these observations. We found no difference in the intercept or the slope of the two regressions (Table 3), which indicates that PE elements have no discernable effect on reproductive senescence.

**TABLE 3.** The rate of decrease of reproductive potential is similar in females PE^+^ and PE^−^ females. Genotype refers to whether females have PEs. Intercepts (Row 1) and slopes (Row 3) are similar in the decline of ovariole number as females age indicating that the genotype has no effect on the initial output or the rate of decay on fertility as age progresses. Df: degrees of freedom.

|  | **Df** | **Sum of the squared differences** | **Mean squared error** | **F-value** | **P-value** |
| --- | --- | --- | --- | --- | --- |
| **Genotype** | 1 | 26 | 25.90 | 0.201 | 0.654 |
| **Age** | 1 | 1,158 | 1158.24 | 8.995 | 0.003 |
| **Genotype × Age** | 1 | 468 | 467.91 | 3.634 | 0.057 |
| **Residuals** | 855 | 110,095 | 128.77 |  |  |

#### Male sterility

We studied male sterility in two ways. First, we dissected the testes of F1 males from crosses between PE‐ and PE+ individuals. At 23°C, no F1 male, regardless of their genotype, showed atrophied testes. All males had motile sperm at this temperature. At 29°C, male sterility was most often observed in individuals produced from the crosses that involved a PE^+^ parent (PE^−^/ PE^+^, PE^+^/ PE^−^, and PE^+^/ PE^+^) than in males with no PEs (PE^−^/ PE^−^; Table 4). The likelihood of obtaining F1 sterile males was slightly higher if the mother carried PEs (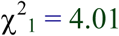, P = 0.045) but not if the father did (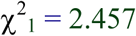, P = 0.117). All LRT tests shown in Table S6.

**TABLE 4.**
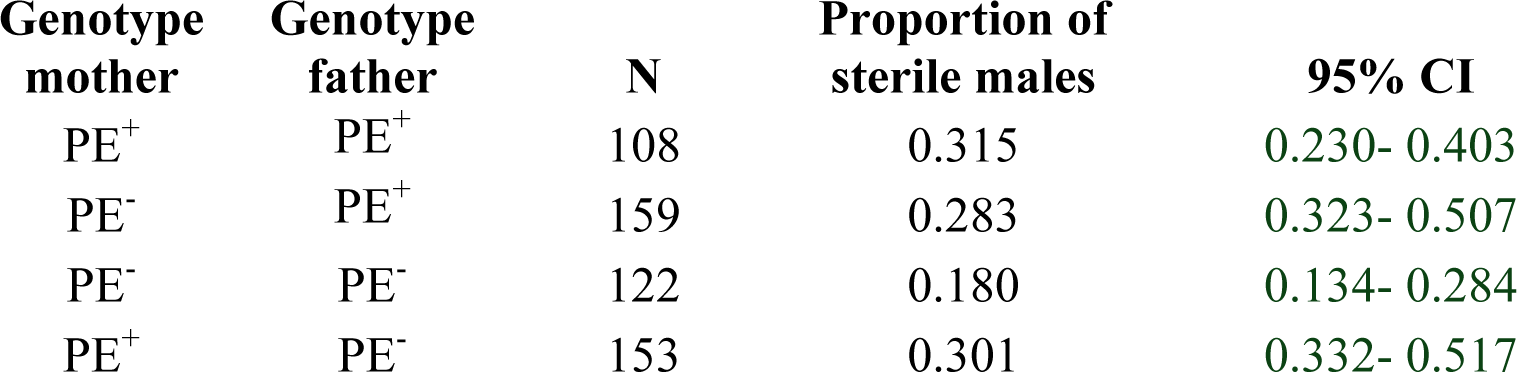
F1 male sterility at 29°C is more likely to occur if any of the parents carry PEs. N is the number of dissected males. 95% CI is the 95% confidence interval of the mean. Table S6 shows the relevant LRTs.

| **Genotype**<br>**mother** | **Genotype**<br>**father** | **N** | **Proportion of**<br>**sterile males** | **95% CI** |
| --- | --- | --- | --- | --- |
| PE<sup>+</sup> | PE<sup>+</sup> | 108 | 0.315 | 0.230- 0.403 |
| PE<sup>-</sup> | PE<sup>+</sup> | 159 | 0.283 | 0.323- 0.507 |
| PE<sup>-</sup> | PE<sup>-</sup> | 122 | 0.180 | 0.134- 0.284 |
| PE<sup>+</sup> | PE<sup>-</sup> | 153 | 0.301 | 0.332- 0.517 |

Second, we scored the fertility of the four different genotypes of males when they mated to PE‐ females. When males were raised either at 23°C or 29°C, we found no differences in the number of progeny produced between genotypes (cross effect: F_3,72_ = 0.639, P = 0.593). Consistent with previous studies (Stanley *et al*. 1980; Matute *et al*. 2009), we found that higher temperatures (i.e., 29°C) reduce male fertility in *D. yakuba* (temperature effect: F_1, 72_ = 4.370, P = 0.040) but no differential effect of temperature on different PE carriers and non-carriers (cross × temperature effect: F_3,32_ = 0.341, P=0.796). Unlike the strong effect of PEs in female fertility (at least at 29°C), the effect of PEs in male fertility is little to non-existent.

## DISCUSSION

P transposable elements (PEs) have rapidly spread across various *Drosophila* species and have invaded their genomes at different rates and geographic locations. In at least two different *Drosophila* species, PEs have reached populations in every continent (Kidwell *et al*. 1977; Kofler *et al*. 2015; Hill *et al*. 2016b). This increase in frequency is surprising because PEs have drastic fitness costs associated with heterotypic matings: F1 progeny from crosses between non-PE containing females and PE containing males are regularly sterile (Kidwell *et al*. 1977; Hill *et al*. 2016b). Here, we show that the genome of some *D. yakuba* individuals now harbor PEs, while its two sister-species (*D. santomea* and *D. teissieri*) do not appear to contain a functional PE. Notably, we find PEs vary in prevalence in the São Tomé population between collection years and although our PCR scans did not detect exons on the continent it is possible that it is present in continental populations at both low population and intra-genome frequencies.

*Drosophila teissieri* (and *D. sechellia* and, to a lesser extent, *D. mauritiana*) poses an interesting case. The detection of PEs using short reads revealed the presence of highly fragmented PEs. This might indicate that PEs were present and active in the past but they are now degenerated. Certainly a larger collection of *D. teissieri* individuals will be needed before this hypothesis can be formally tested.

*Drosophila yakuba* is the third species in the *melanogaster* group found to be infected by PEs, and unlike other species, PEs are still actively segregating at a low frequency. Our findings have three broad implications*: i)* that the increase in frequency of PEs is not always monotonic after an invasion occurs *ii)* they indicate that PEs in *D*. *yakuba* cause a much milder hybrid dysgenesis syndrome than that observed in *D. simulans* and *D. melanogaster*, and *iii*) pose the possibility of transmission of PEs through hybridization and subsequent introgression.

### Variable frequency of PEs across time points

*Drosophila yakuba* represents the latest case of an invasion of PEs in natural populations. Of all the known invasions, it also represents the only case in which a decrease in PE frequency over time has been noted in natural populations. In both *D. simulans* and *D. melanogaster*, the invasion of PEs was discovered when it was widely distributed across worldwide populations. These three invasions represent a natural system in which to explore TE spread and the evolution of repressive systems to counter it.

A similar approach has used artificial invasions of PEs to understand how fast they occur. Kofler et al. (Kofler *et al*. 2018) studied the genome invasion by PEs in *D. simulans* and found that the process had two stages. First, and rapidly after the PE introduction, PEs increased in frequency, especially at high temperatures. In the second stage, PE frequency plateaued at a high frequency throughout the population but did not fix. Notably, the number of copies of the PE per genome in this *D. simulans* invasion were similar to those observed in *D. melanogaster* (Figure 1B in Kofler et al. 2018 and Figure 1). The magnitude of the natural invasion in *D. yakuba* here reported is much smaller and more akin to the rate of transposition observed in the experimental invasion of *D. simulans* at lower temperatures.

Surprisingly, in contrast to the allelic and geographic spread of PEs witnessed in *D. melanogaster* and *D. simulans*, in *D. yakuba* we see a rapid increase (18%) in frequency between 2013 and 2015, subsequently followed by a pronounced drop (2%) in 2018. Seasonal variation in genetic frequencies over time seems to be a common phenomenon in nature (Bergland *et al*. 2014). Longitudinal studies of *Drosophila* populations in temperate regions have found temporal variation in multiple genomic regions which have been hypothesized to be related to seasonal fitness variation related to temperature and humidity. The range of *D. yakuba*, and São Tomé in particular, are tropical environments and do not experience changes in temperature as large as temperate areas of the globe but they still show environmental yearly cycles. A systematic longitudinal collection will be required to answer the nature and amplitude of the observed PE frequency decline.

A second genetic element that shows variation in frequency across years in *D. yakuba* from São Tomé is *Wolbachia*. Between 2001 and 2009, the frequency of *Wolbachia* in *D. yakuba* experienced an increase from 25% to 75%. Between 2009 and 2015, there was no increase in infection frequency (Cooper *et al*. 2017a). A finer temporal scale sampling is needed to resolved whether this variation is related to seasonality or corresponds to longer cycles. Regardless of the actual timescale, whether the variation is explained by seasonal cycles or the amplitude of the period is longer, the variability in *Wolbachia* and PE elements in species found in São Tomé suggest that seasonal temporal studies need to be conducted in tropical populations as well as temperate ones.

Although the mechanism that led to such a drastic decline in infected individuals between 2015 and 2018 is unknown, there are two possibilities. *Wolbachia* has been hypothesized to prevent virus infection; if PEs are transmitted by viruses (and idea that remains highly speculative) the increase in frequency of *Wolbachia* could be responsible for the decrease in PE frequency. Such protective effects have been reported in mosquitoes (van den Hurk *et al*. 2012; Lee *et al*. 2013; Johnson 2015; Aliota *et al*. 2016) and *Drosophila* (Hedges *et al*. 2008; Osborne *et al*. 2009, 2012; Martinez *et al*. 2014; Shi *et al*. 2018). In the latter case, the magnitude of the protection is highly contingent on the *Wolbachia* strain and the genotype host suggesting a strong genetic interaction (Longdon *et al*. 2012; Martinez *et al*. 2017). This potential protective effect of *Wolbachia* could explain a decrease in horizontal gene transfer of PEs into *D. yakuba* but not a decrease in frequency of PEs after they had invaded. Given the lack of evidence for the involvement of viruses on the transmission of PEs, this possibility should be considered speculative. An additional possibility is that São Tomé has experienced an increase in temperature and such conditions lead to the decrease of the frequency of PEs. If *D. yakuba* flies carrying PEs show a decreased fitness at higher temperatures, then PEs might be expected to decrease in frequency. Yet, tropical populations of *D. simulans* and *D. melanogaster* have also seen an increase in their PE frequency, which would argue against this possibility. Additionally, PEs increase in frequency much more rapidly in synthetic populations of *D. simulans* at higher temperatures (Kofler et al. 2018). It is worth nothing that of the three species from the *melanogaster* subgroup in which the PE has been found, *D. yakuba* is the most sensitive to high temperatures (Stanley *et al*. 1980; Matute *et al*. 2009). The interaction between different environments, such as temperature differences, and the rate of spread of PEs remains mostly unexplored (but see (Kofler *et al*. 2018)).

### Mild hybrid dysgenesis in *D. yakuba*

We found that *D. yakuba* is polymorphic for the presence of PEs. We only found evidence of PEs in the island of São Tomé where they appear to have invaded between 2013 and 2015 but found no evidence for PEs in the African continent or the islands of Principe and Bioko using our PCR scans. This result is puzzling as population structure between different *D. yakuba* populations is low (Comeault *et al*. 2016), thus we would expect PEs to spread as the deleterious effects of PEs are limited compared to those in *D. melanogaster* and *D. simulans*. This recent invasion of PEs in *D. yakuba*, and more precisely of the populations in São Tomé, represents a unique opportunity to study a transposable element invasion into a naïve genome and witness how the genome adapts throughout time (Kofler *et al*. 2018).

In *Drosophila,* the deleterious phenotypes of HD tend to manifest in the F1 progeny of PE^−^ females and PE^+^ males. We assessed the existence of four possible phenotypes associated with PE-induced hybrid dysgenesis: *i*) presence of atrophied ovaries, *ii*) reduced number of ovarioles, *iii*) early onset of reproductive senescence, and *iv*) reduced male fertility. These defects are all associated to hybrid dysgenesis in *D. melanogaster* (Kidwell *et al*. 1977; Khurana *et al*. 2011). In *D. simulans*, hybrid dysgenesis is known to cause atrophied ovaries but defects *ii*-*iv* have not been explored in relation to PEs. Of the traits we measured, we found that the only manifestation of hybrid dysgenesis in *D. yakuba* is an increase in the number of atrophied ovaries in all crosses that involved individuals carrying full PEs, but only at 29°C, and most frequently in F1 females from the cross ♀PE^−^/♂PE^+^. This mild manifestation of hybrid dysgenesis might be associated with the recent invasion of the PEs in *D. yakuba*, resulting in low PE copy number (Nuzhdin 2000). Currently, we have no information as the number of PE copies in each *D. yakuba* genome, nor its distribution in the genome, but future assemblies with long reads should be able to address these questions.

Surprisingly, we see an increase in ovariole number in F1s that result from PE+ *D. yakuba* females being crossed to PE^−^ *D. yakuba* males. This result is intriguing as it suggests a potential fitness benefit to PEs into novel genomic backgrounds, a result that has not been previously described. Comparisons of PE^+^ and PE^−^ *D. yakuba* isolines show no differences between the parentals used to obtain the F1s, therefore maternal/paternal differences in fecundity are unlikely to explain this result. If an increase of ovariole number in a PE+/PE‐ cross is only seen in *D. yakuba* or also occurs in *D. melanogaster* and *D. simulans* remains to be tested. If this pattern holds across distinct species, it can provide an explanatory driving force behind the rapid expansion of PEs across worldwide populations, in terms of coupling a fitness advantage with innate TE’s self-replicating nature.

### Transmission of PEs between species

PEs are thought to have originated in the Neotropical *willinstoni* group, a group 50-60 million years separated from the *melanogaster* lineage split (Throckmorton 1975; Beverley and Wilson 1984). Our findings of PEs recently incorporating into the *D. yakuba’* genome, deepens the puzzle of how PEs move across species boundaries. None of the species from the *melanogaster* species subgroup can hybridize with species from the *willinstoni* group. In the case of *D. melanogaster* and *D. simulans,* crosses produce hybrid progeny from the sex of the mother and in most cases the progeny is sterile. One exception is crosses between *D. melanogaster* In(1)AB females which can produce fertile female F1s when crossed with the *D. simulans* strain C167.4 (Davis *et al*. 1996). These hybrid viability rescue mutations in the genes *Lhr* and *Hmr* seem to segregate at very low frequency in nature and are unlikely to constitute a bridge for gene transfer. *Drosophila melanogaster* and *D. yakuba* also can produce viable hybrids, only when behavioral isolation is circumvented, but the resulting hybrids are sterile in all cases (Sánchez and Santamaria 1997). Similarly, *D. simulans* females and *D. yakuba* males produce viable, yet sterile, female offspring (Orr 1993; Turissini *et al*. 2018).

The high level of similarity of the PE sequences across all species, and the longitudinal data collected for *D melanogaster*, *D. simulans*, and *D. yakuba*, are consistent with a recent transmission of this genetic element across species of *Drosophila*. Since hybridization does not seem to be the mode of transmission of PEs (see above), the possibility of horizontal gene transfer seems more likely. Horizontal gene transfer (HGT) has been hypothesized as a major mechanism of distribution across various transposable elements (Keeling and Palmer 2008; Schaack *et al*. 2010). The leading hypothesis regarding interspecific HGT of PEs (in particular the transfer of PEs from *D. willinstoni* into *D. melanogaster*) argues that the mouthpieces of the mite *Procteolaelaps regalis*, who feeds on *Drosophila* eggs, might function as a micro-injection device and might facilitate DNA transfer between embryos. Support for this hypothesis stems from the presence of PEs in both mites and flies, and the ecological overlap of *D. melanogaster*, *D. willistoni* and *P. regalis* (Houck et al. 1991). Yet this mechanism has not been directly demonstrated (Houck *et al*. 1991; Engels 1992). Other non-mite vectors are also possible, but have been less explored. For example, TEs can also be transmitted through viruses (Loreto *et al*. 2008) but the relevance of viruses on HGT in *Drosophila* is largely underexplored. As with other cases of HGT, the precise mechanism through which TEs have successfully spread across the majority of higher order Eukaryotes, making up large portions of their genome, is not well understood, and further exploration is required to understand the role of HGT in eukaryotic evolution (Fedoroff 2012).

Notably, *D. yakuba* hybridizes with two other species and produce stable hybrid zones in the islands of São Tomé and Bioko (*D. santomea* (Llopart *et al*. 2009; Comeault *et al*. 2016) and *D. teissieri* (Cooper *et al*. 2017b) respectively). As of 2018 neither species contains active PEs. This raises the question of why PEs have not crossed from *D. yakuba* into *D. santomea* nor *D. teissieri*. Areas of secondary contact, as well as laboratory crosses, will reveal whether PEs are prone to be transferred through introgression or whether their interspecific transfer is penalized at a greater rate between species than within (Waugh O’Neill *et al*. 1998; Labrador *et al*. 1999). Another naturally hybridizing species that might serve as potential system through which to explore the role of hybridization in the transfer of PEs occurs in the Seychelles archipelago. *Drosophila simulans* and *D. sechellia* hybridize in the central islands of the archipelago where human density is the highest (Matute and Ayroles 2014). PEs have been found in *D. simulans* but not in *D. sechellia,* similar to the observations of the presence of PEs in *D. yakuba* but not in its sibling species. These hybrid zones should be explored to assess potential expansions or limitations of PEs across species boundaries through hybridization and introgression.

## Conclusions and future directions

A precise quantification of the frequency of PEs, and TEs in general, across different species is still in its infancy (Watson and Demuth 2013; Serrato-Capuchina and Matute 2018). Even though the phenomenon of hybrid dysgenesis has been rigorously characterized in *D. melanogaster* (Kelleher 2016), the discovery of PEs in other *Drosophila* species allows us to understand how these elements behave in different genetic backgrounds. In the case of *D. yakuba*, PEs cause a hybrid dysgenesis phenomenon which is much milder than in *D. melanogaster* and almost exclusively manifests at 29°C. We do not know whether this comparatively minor dysgenesis is caused by the recency of the invasion (which would lead to potentially few copies of the PE), or the genetic background of *D. yakuba*. In any case, these results suggest that the invasion of PEs in a genome does not have the deterministic outcome of hybrid dysgenesis, and instead these deleterious effects might be modulated by the timing of the invasion and/or genetic background of the invaded species. Other reports have found the same element in mites which suggest the PE is present in other arthropods (Houck *et al*. 1991). The results shown here strongly suggest that a population level assessment of the presence of PEs in a wide variety of taxa will be needed before we understand the precise taxonomic distribution and effects of PEs.

## ACKNOWLEDGEMENTS

We would like to thank, A.A. Comeault, D.A. Turissini, G. Bates for assistance in the field. C.J. Jones, B.S. Cooper, and members of the Matute lab, gave us helpful feedback on the manuscript. This work was supported by NIH award R01GM121750 to D.R.M‥ The authors declare no conflicts of interest.

